# Use-dependent regulation of the axonal action potential in parvalbumin-expressing interneurons

**DOI:** 10.64898/2026.02.07.704588

**Authors:** Sophie R. Liebergall, Ethan M. Goldberg

**Affiliations:** Department of Neuroscience, The University of Pennsylvania Perelman School of Medicine, Philadelphia, PA, USA; Neuroscience Graduate Group, The University of Pennsylvania Perelman School of Medicine, Philadelphia, PA, USA; Medical Scientist Training Program, The University of Pennsylvania Perelman School of Medicine, Philadelphia, PA, USA; Department of Neurology, The University of Pennsylvania Perelman School of Medicine, Philadelphia, PA, USA; Division of Neurology, The Children’s Hospital of Philadelphia, Philadelphia, PA, USA

**Keywords:** Action potential, axon, interneuron, seizure, epilepsy

## Abstract

The action potential (AP) is thought to be generated at the axon initial segment and to faithfully propagate along the axon. However, data from both invertebrate and mammalian systems show that the axon is an underappreciated locus of activity modulation and neuronal computation. We assessed axonal AP propagation in neocortical parvalbumin-expressing interneurons (PVINs) during prolonged, high-frequency activity through paired whole-cell somatic and axon-attached patch clamp recordings in acute brain slices from mouse and human. We found that PV-IN axonal AP propagation remains robust during prolonged activity at moderate frequencies, such as during the entrainment to PV-IN firing patterns recorded in awake, behaving mice *in vivo*. However, prolonged, high-frequency activity during evoked trains of APs and during seizure-like events resulted in changes in the waveform of the axonal (but not somatic) AP, at least in part due to intrinsic use-dependent mechanisms. This use-dependent decrement in the axonal AP waveform is associated with decreases in calcium influx at PV-IN boutons and subsequent PV-IN-mediated synaptic transmission, indicating this phenomenon may lead to a dissociation between somatic and axonal excitability that could shape PV-IN contributions to circuit dynamics during periods of high activity.

## Introduction

The dominant view of neuronal signaling asserts that action potentials (APs) are generated at the axon initial segment (AIS) and faithfully propagate down the axon to trigger neurotransmitter release at the presynaptic terminal. However, work in both invertebrate and vertebrate systems shows that the axon is not simply a cable for the digital transmission of somatic/AIS signals, but instead constitutes a distinct site for analog regulation (1–5). Diversity in the anatomical properties of axons across cell types is so profound that axonal morphology has historically served as a major characteristic by which cell type has been defined (6, 7). However, assessing the physiological properties of axons has been hampered by the technical challenges of directly recording from such thin, complex, and spatially extended structures (5). The axons of cerebral cortex parvalbumin-expressing interneurons (PV-INs) may be particularly susceptible to axon-specific regulation of the axonal AP for several reasons: PV-IN axons support high-frequency firing, are only intermittently myelinated, exhibit extensive branching, and are interrupted by *en passant* boutons (8–11).

Given that PV-INs control the activity of glutamatergic excitatory neurons, it has been hypothesized that failure of PV-IN inhibition and subsequent runaway excitation is a key mechanism of focal-onset seizure, which is a brain state defined by increased, hypersynchronous neural activity (12– 15). However, numerous studies in both human patients and animal models have documented *increased* firing of PV-INs prior to and at seizure initiation (16–21). Studies reporting PV-IN firing during the onset and propagation of seizures or seizure-like events have primarily employed techniques such as intra/extracellular somatic electrophysiology and 2-photon Ca^2+^ imaging, which are limited to recording of PV-IN firing at the soma (16, 21–26). We hypothesized that decreased fidelity of AP propagation along PV-IN axons could serve as a mechanism contributing to failure of inhibitory restraint.

In this study, we directly assessed axonal AP propagation in neocortical PV-INs during prolonged epochs of high-frequency activity. We found that axonal AP propagation remains robust even during prolonged activity at moderate frequencies (<100 Hz), such as during PV-IN firing patterns observed in awake, behaving mice *in vivo*. However, prolonged high-frequency activity (>100 Hz)—observed both during evoked AP trains and in an *ex vivo* model of seizure-like events—led to profound use-dependent changes in the axonal AP waveform. This modulation involved cell-intrinsic, use-dependent mechanisms, could be well described by a sum of exponential decays, and were observed in both mouse and human fast-spiking interneurons.

## Materials and Methods

### Experimental animals

All procedures and experiments were approved by the Institutional Animal Care and Use Committee at the Children’s Hospital of Philadelphia and were conducted in accordance with the ethical guidelines of the National Institute of Health. All studies used both male and female mice in equal proportions. Mouse strains used in this study included: PV-Cre mice (B6.129P2Pvalbtm1(cre)Arbr/J; RRID: IMSR_JAX:017320), Ai14 (tdT) mice (B6.Cg-Gt(ROSA)26Sortm14(CAGtdTomato)Hze/J; RRID: IMSR_JAX:007914), and WT 129S6/SvEvTac (Taconic; RRID: IMSR_TAC:129SVE). Mice homozygous for PV-Cre/Ai14 were crossed to WT 129S6/SvEvTac mice to generate experimental animals heterzygous for PV-Cre/Ai14.

### Human surgical specimens

All experiments using human tissue were done in accordance with the Children’s Hospital of Philadelphia Institutional Review Board’s approval (IRB 15-012226). Tissue was retrieved in the operating room in pre-oxygenated ice-cold ACSF and prepared as for the mouse tissue. Cortical fast-spiking interneurons (presumably parvalbumin positive) were identified by their fast-spiking physiology (steady-state firing frequency >100 Hz), and nonpyramidal somatic morphology.

### Acute slice preparation

To prepare acute brain slices for electrophysiology experiments, mice aged P16-21 were anesthetized with isoflurane and the brain was transferred to ice cold artificial cerebral spinal fluid (ACSF) containing (in mM): NaCl, 87; sucrose, 75; KCl, 2.5; CaCl_2_, 1.0; MgSO_4_, 6.0; NaHCO_3_, 26; NaH_2_PO_4_, 1.25; glucose, 10, and equilibrated with 95% O2 and 5% CO_2_. The brain was sliced at a thickness of 300 µm using a Leica VT-1200S vibratome. Slices were allowed to recover in ACSF for 30 mins at 32°C, then maintained at room temperature for up to 4 hours before recording. For recordings, slices were transferred to the recording chamber and continuously perfused at a rate of 3 mL/min and temperature of 31-32° C with a recording solution that contained, in mM: NaCl, 125; KCl, 2.5; CaCl_2_, 2.0; MgSO_4_, 1.0; NaHCO_3_, 26; NaH_2_PO_4_, 1.25; glucose, 10. For recordings in high (10 mM) K^+^, the recording solution contained, in mM: NaCl, 117.5; KCl, 10; CaCl_2_, 2.0; MgSO_4_, 1.0; NaHCO_3_, 26; NaH_2_PO_4_, 1.25; glucose, 10. For human recordings, human surgical specimens were transferred to the same cutting ACSF that is used for mouse recordings and were immediately transported to the lab (<10 min) and sectioned as above. Recordings were performed in slices of healthy neocortex that was removed in the process of gaining access to structural pathology (*N* = 3: 8 year old, lateral temporal cortex, focal cortical dysplasia; 5 year old, frontal cortex, focal cortical dysplasia; 6 month old, lateral temporal cortex, focal cortical dysplasia). Recordings were performed as in mouse tissue, with the exception that slices were maintained at room temperature for up to 24 hours.

### Paired whole-cell somatic and axon-attached acute brain slice recordings

All recordings were performed using a commercial 2-photon microscope (Bruker) equipped with a Leica objective (25X, 0.95 NA), two GaAsP photocathode photomultiplier tubes (H10770PA-40, Hamamatsu), and a substage detector for simultaneous laser-scanning Dodt contrast. A femtosecond-pulsed Ti:sapphire laser (Insight X3^+^, Spectra Physics) was tuned to λ = 950 nm for fluorescence excitation. PV-INs were identified by red fluorescence expression visualized with 2-photon imaging. Wholecell recordings were obtained from Layer 2/3 PV-INs in somatosensory cortex. Patch pipettes were pulled from borosilicate glass using a P-1000 puller (Sutter Instruments). Somatic whole cell pipettes with a resistance of 3–5 MO were filled with intracellular solution containing (in mM): K-gluconate, 130; KCl, 6.3; EGTA, 0.5; MgCl_2_, 1.0; HEPES, 10; Mg-ATP, 4.0; Na-GTP, 0.3; 100 µM Alexa Fluor 488 hydrazide (Thermo Fischer); pH was adjusted to 7.30 with KOH and osmolarity was adjusted to 290 mOsm. In paired somatic recording of unitary synaptic transmission, recording pipettes used for the post-synaptic cell were filled with an intracellular solution containing K-gluconate, 65; KCl, 65; MgCl_2_, 2; HEPES, 10; EGTA, 0.5; phosphocreatine-Tris, 10; Mg-ATP, 4.0; Na-GTP, 0.3 (E_Cl_ = −17 mV to enable larger signal to noise ratio for recordings of unitary inhibitory post-synaptic currents). For axon-attached recordings, patch pipettes with a resistance of 5-8 MO were filled with recording ACSF. Somata were first patched in the whole-cell configuration using the laser-scanning Dodt contrast image and the Alexa Fluor 488 dye was allowed to diffuse for 20-30 mins until the distal axon could be visualized using the 2-photon image. Distal axon was distinguished from dendrites on the basis of smaller diameter and the presence of extensive wide-angle branching and abundant varicosities *en passant* boutons). Gentle positive pressure was applied to the axon-attached recording pipette and an intact branch of the distal axon was approached perpendicularly. Positive pressure was released and a loose seal (generally 10-25 MO) was formed. Signals were sampled at 100 kHz with a MultiClamp 700B amplifier (Molecular Devices), filtered at 10 kHz, digitized using a DigiData 1550A, and acquired using pClamp10 software. Recordings were discarded if the cell had a somatic resting membrane potential was more depolarized than −50 mV, if somatic access resistance was > 25 MO, or if the axon-attached signal was < 10 pA after an attempt at forming a loose seal. Series resistance compensation (bridge balance) was applied at the soma pipette. We did not correct for the liquid-liquid junction potential. The command potential of the voltage for the axon-attached recording in voltage-clamp was set such that the amplifier injected a current of 0 pA, so as to avoid influencing the membrane potential of the recorded axon (27). At the conclusion of each recording, a tiled 2-photon *z-*stack of the morphology of the recorded cell was obtained.

### Simulated normal physiology in vivo spike pattern

The *in vivo* spike pattern delivered as a pulse train to cells in Figure 4 was based on data from the Allen Institute MindScope Program’s Allen Brain Observatory Neuropixels Visual Behavior dataset (2022, available from https://portal.brain-map.org/circuits-behavior/visual-behavior-neuropixels). Spike pattern extraction and generation of the stimulus file were performed using custom Python code. A recording session from a Pvalb-IRES-Cre/wt;Ai32(RCL-ChR2(H134R)\_EYFP)/wt mouse was selected where optotagging was performed at the conclusion of the session with an implanted blue laser (maximum power at fiber tip = 35 mW). Spikes of identified units were aligned to the delivery of 10 ms light pulses, and cells that fired at >100 Hz in response to the light pulses and displayed a narrow mean waveform consistent with a fast-spiking IN were selected. Of the selected units, a single unit that displayed an average firing rate (>20 Hz) was chosen, and spike times were extracted from a 5 min period when the cell showed peak activity (when the mouse was passively viewing drifting gratings). A .atf file containing brief depolarizing pulses at times corresponding with the *in vivo* PV-IN spike times was generated using custom code adapted from the pyABF package (S.W. Harden, pyABF 2.3.5., available from https://pypi.org/project/pyabf).

### Electrophysiological data analysis

All analysis was performed with custom code in MATLAB 2023b (MathWorks). To determine the fidelity of axonal AP propagation during paired whole-cell somatic and axon-attached recordings, first the locations and amplitudes of soma APs were determined using the findpeaks function, where an AP was defined as a spike that reached a threshold of at least −10 mV and peak prominence of at least 20 mV. For seizure-like event recordings, in which there is larger variability in the height of somatic APs, a threshold of −20 mV was used. Next, the locations and magnitudes of the AP current peaks in the axonal recording were determined by (1) creating a 1 ms window centered on the peak of each somatic AP, (2) finding the minimum value in the axonal current signal to determine the location of the axonal AP peak, (3) calculating the mean value of the axonal current signal 0.5-1 ms before the axonal AP peak to determine the baseline axonal current, and (4) calculating the difference between the baseline and peak axonal AP current to determine the magnitude of the inward axon AP peak (in pA). Soma-axon latency was defined as the difference in the time of the axon AP peak current and the time of the soma AP peak voltage.

### Morphology reconstructions

Tiled 2-photon *z*-stacks of recorded cells, in which the 2P green channel contained the morphology of the recorded cell from the Alexa Fluor 488 fill, the 2P red channel contained the tdT (or mCherry) expression in the target population, and the Dodt channel contained information about the locations of the recording pipettes, were combined into single 3-channel images using the Neurolucida 360 software (MBF Bioscience). The Neurolucida 360 software was used to manually trace the morphology of the soma, axon, and dendrites of the recorded cells in 3-D. After the reconstruction was complete, the Neurolucida Explorer software (MBF Bioscience) was used to calculate the recording distance (the 3-D length of the path from the axon hillock to the axonal recording site) and the branch order (the number of axonal branch points between the axon hillock and the axonal recording site) for each recording.

### Bouton Ca^2+^ imaging

The same microscope that was used for slice electrophysiology experiments (see above) was also employed for bouton Ca^2+^ imaging experiments. Neurons were patched at the soma in the whole-cell configuration using 3-5 MO pipettes filled with intracellular solution containing (in mM): K-gluconate, 126; KCl, 4; HEPES, 10; MgATP, 4.0; Na_2_-phosphocreatine, 10; Fluo-5F, 0.1 (F14221, Thermo Fischer), Alexa Fluor 594 Hydrazide 0.1 (Thermo Fischer); pH was adjusted to 7.30 with KOH and osmolarity was adjusted to 290 mOsm. Cells were allowed to fill for 20-30 mins before imaging. Red fluorescence was used to locate boutons on the PV-IN distal axon. Green fluorescence (change in Ca^2+^ concentration) and red fluorescence (structural image) were measured while scanning with a resolution of 125×125 pixels at 75 Hz. Initiation of the bouton imaging scan was synchronized with the initiation of the delivery of 100 Hz pulses at the somatic recording pipette using the PrairieView imaging software (Bruker). Analysis of bouton Ca^2+^ imaging was performed using custom code in Matlab 2023b (Mathworks). First, 15×15 pixel ROIs centered on each bouton were manually selected from each field of view using the mean image from the recording. Next, average red and green fluorescence values were extracted from each ROI at each frame. ROIs were excluded from analysis if there was significant *xy* or *z* movement during the recording evident in the structural (red) channel. We did not observe any photobleaching in the red channel during recordings. dG/R values were calculated using the formula:

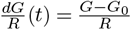

where *G* is the green fluorescence at time *t, G*_0_ is the baseline fluorescence, and *R* is the red fluorescence at time *t* (28). Individual traces were smoothed with a median filter (`medfilt1`) for visualization.

### Sum of exponential decays model

Use-dependent decreases in axon AP amplitude were modeled in Matlab 2023b (MathWorks) using a sum of exponential decays defined by the following equation:

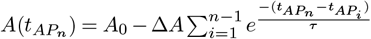

where 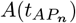 is the peak of the n^th^ AP, *A*_0_ is the baseline offset, ∆*A* is the scaling factor, *τ* is the decay time constant, 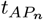 is the time of the n^th^ AP, and 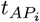 is the time of the i^th^ AP. Axon AP peak values (normalized to the mean of the first 5 APs) for each recording were fit using non-linear least-squares curve fitting (`lsqnonlin`), initialized with values of *A*_0_ = maximum *A*_*t*_ value in recording, 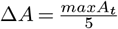, and *τ* = 10 ms. Decay time constants *τ* were estimated within the following constraints: *τ* > 0 ms and *τ* < 200 ms. 2 recordings reached the fitting parameter boundary limit and were excluded from subsequent analyses.

### Quantification and Statistical Analysis

*N* values for all experiments, definitions of center, and definitions of dispersion values are indicated in the figure legends. Efforts were made to display individual data points whenever possible. Statistical comparisons were not required for most experiments, but when necessary were detailed in the figure legend.

## Results

### Use-dependent modulation of PV-IN axonal AP propagation

Previous studies have shown that PV-IN axonal AP propagation is reliable during brief trains of APs stimulated by somatic current injections *in vitro* (10, 11, 29). However, PV-INs can sustain high firing rates (>50 Hz) for seconds to minutes *in vivo* (30–33), but whether APs faithfully propagate to the synapse under such conditions is unknown. To assess the impact of such prolonged high-frequency firing on axonal AP propagation, we performed paired whole-cell somatic and axon-attached recordings in neocortical PV-INs from layer 2/3 of primary whisker somatosensory (“barrel”) neocortex under 2-photon guidance in acute brain slices from P16-21 wild type (WT) mice in response to application of brief depolarizing current pulses at the soma at varying frequencies (Figs. 1-2). Consistent with previous reports, stimulation at 50 Hz did not lead to changes in the axonal AP waveform on relatively short timescales (hundreds of milliseconds; Fig 2B) (34). However, high-frequency stimulation at 100 Hz (or higher) led to marked changes in the axonal AP waveform within seconds (Fig. 2C-H). These use-dependent changes in the axonal AP waveform outpaced corresponding changes in both the voltage trace of the somatic AP as well as the max dV/dt of the somatic waveform (Fig. S1). For each recording, we obtained a 2-photon *Z*-stack of the cell and reconstructed the axonal path from the axon hillock to the recording site to quantify the distance (range, 48.5 to 455.1 µM) and branch order (range, 1 to 15) of the recording site (Figs. 1, S3). Notably, the magnitude of the change in the axonal AP waveform is correlated with the distance along the axonal arbor and the number of branch points that an AP must traverse to reach the recording site (Fig. S3A-B). We also assessed axonal AP propagation while stimulating cells using a standard square wave current steps protocol, in which a 600 ms current step was applied every 3 seconds, starting at −100 pA and increasing by 25 pA steps (Fig. S2). We observed the same phenomenon as depicted in Fig. 2, with PV-INs showing faithful axonal AP propagation at lower current injections and progressive changes in the axonal AP waveform during high current injections (that resulted in higher-frequency firing) (Fig. S2B-D). These apparent decrements in the axonal AP waveform are likely not due to a use- or time-dependent change in the quality of the axon-attached recording configuration, as full-height axonal AP currents were recorded when the cell was again stimulated after a brief “rest” period (Fig. S2E).

**Fig. 1.**
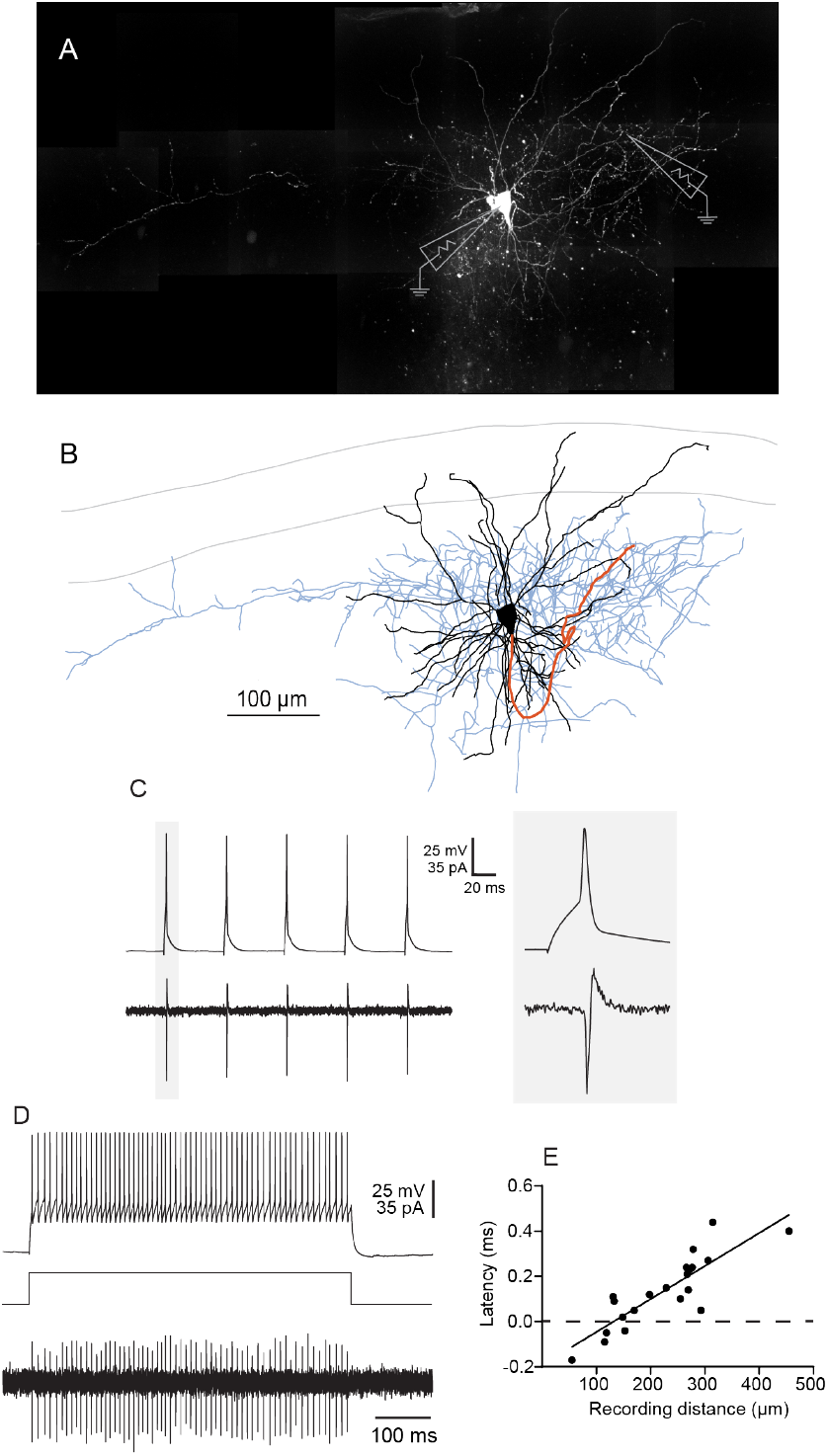
Paired whole-cell somatic and axon-attached recordings in neocortical PV-INs. (A-B) Example (A) 2-photon *z-*stack and (B) morphologic reconstruction of a PV-IN in Layer 2/3 of mouse primary somatosensory cortex (soma and dendrites in black, axon in blue). The path from the axon hillock to the axonal recording site is indicated in orange. (C-D) Example somatic whole-cell voltage traces (top) and axon-attached current traces (bottom) during (C) brief depolarizing current pulses and (D) a depolarizing current step injected through the somatic pipette. (E) Plot of latency from the somatic to the axonal AP peak vs. the distance from the axon hillock to the axonal recording site (*n* = 32 cells from *N* = 23 mice).

**Fig. 2.**
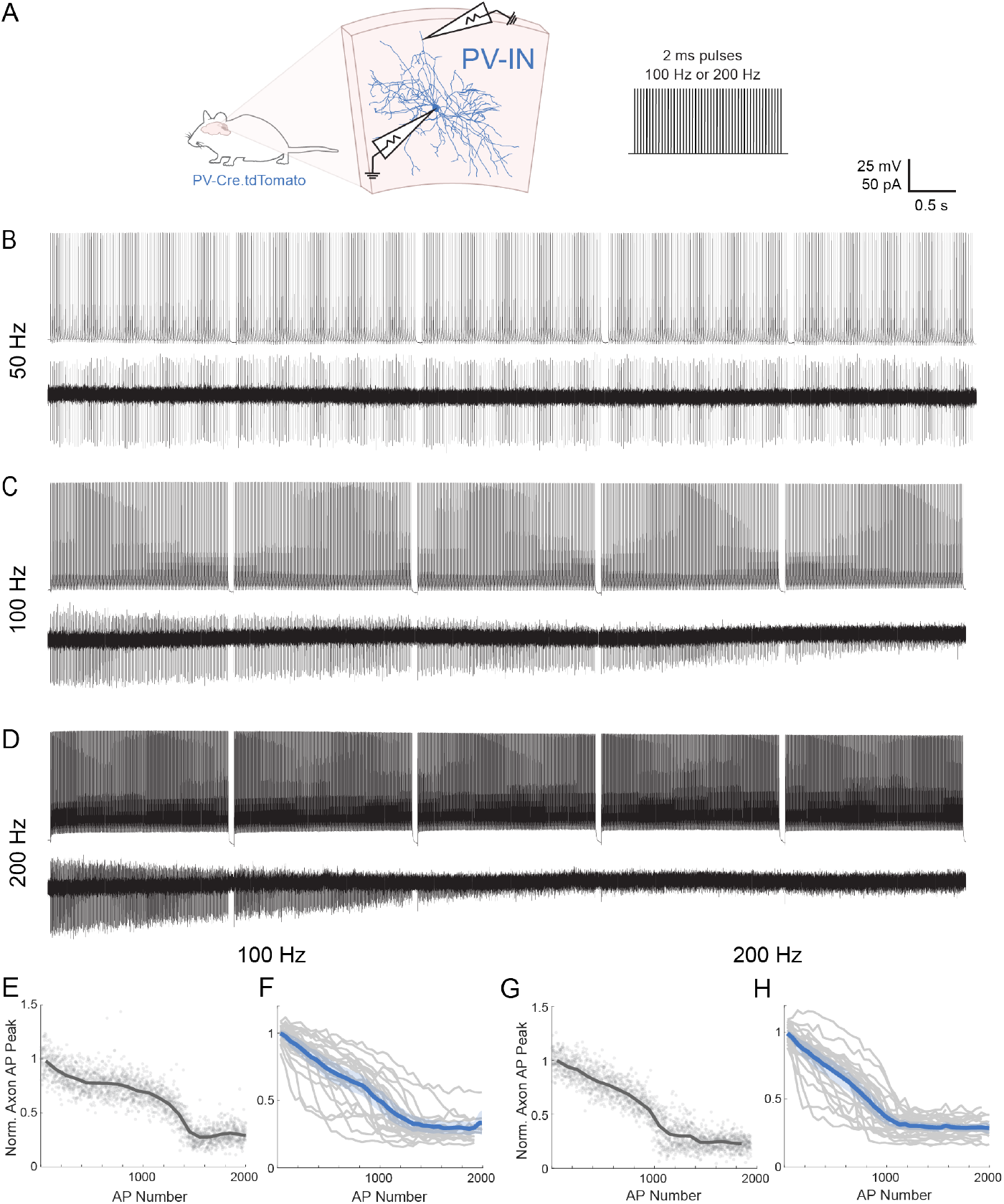
PV-INs display use-dependent changes in the axonal AP waveform during brief depolarizing pulses. (A) Experimental schematic for paired whole-cell somatic and axon-attached recordings in neocortical PV-INs during brief depolarizing pulses. (B-D) Examples of simultaneous whole-cell somatic voltage (top) and axon-attached current (bottom) traces during brief depolarizing pulses delivered at the soma pipette at (B) 50 Hz, (C) 100 Hz, and (D) 200 Hz. (E,G) Plot of axonal AP peak (normalized to the mean of the first 5 APs) vs. AP number for an example cell at (E) 100 Hz and (G) 200 Hz. Gray points indicate individual APs. The dark gray line indicates the mean across bins of 20 APs. (F,H) Summary data of axonal AP peak (normalized to the mean of the first 5 APs) vs. AP number at (F) 100 Hz and (H) 200 Hz. Gray lines indicate individual cells, the dark blue line indicates the mean across cells, and the shaded blue region indicates 95% confidence intervals (*n* = 25 cells from *N* = 14 mice).

Following prolonged trains of soma/AIS-generated APs, neocortical and hippocampal PV-INs in acute brain slices have been reported to enter a persistent firing (or “barrage”) mode in which prolonged trains of APs are spontaneously generated at a site distal to the AIS (35, 36). These axon-generated APs are then propagated retroaxonally, and can be recorded at the soma as either full-height APs or as spikelets (36). During prolonged somatic stimulation of PV-INs, we occasionally observed spontaneous persistent firing (Fig. S4). Given the difference in the latency from soma to axon AP peak between APs generated during somatic stimulation and

APs occurring during persistent firing, we confirmed that APs recorded during persistent firing are indeed generated at a site in the axon distal to the AIS. We examined recordings in which the PV-IN displayed spontaneous persistent firing of distal axon-generated APs and found that, in most recordings, there were progressive changes in the waveform of these distal axon-generated axonal APs (Fig. S4A). Of note, we also observed instances of persistent firing in which propagation from the presumptive distal site of AP generation to the recording site in the axon was of higher fidelity than to the soma (Fig. S4B). This suggests that decreases in axonal AP propagation fidelity are not solely due to the introduction of a recording pipette to the distal axon but can occur between sites along the axon at which no recording pipette is present.

To assess whether the identified use-dependent modulation of the axonal AP is specific to murine PV-INs, we also performed paired whole-cell somatic and axon-attached recordings in fast-spiking INs from grossly normal neocortex obtained from the margins of brain tissue obtained from pediatric patients undergoing surgical resection for treatment of epilepsy (Fig. 3). We recorded 3 human aspiny fast-spiking interneurons with post-hoc morphology consistent with putative basket cells (Fig. 3). In 3/3 cells, we similarly observed use-dependent decrements in the amplitude of the axonal AP waveform (Fig. 3C-D). These findings suggest that use-dependent modulation of the PV-IN axonal AP occur across mammalian species.

**Fig. 3.**
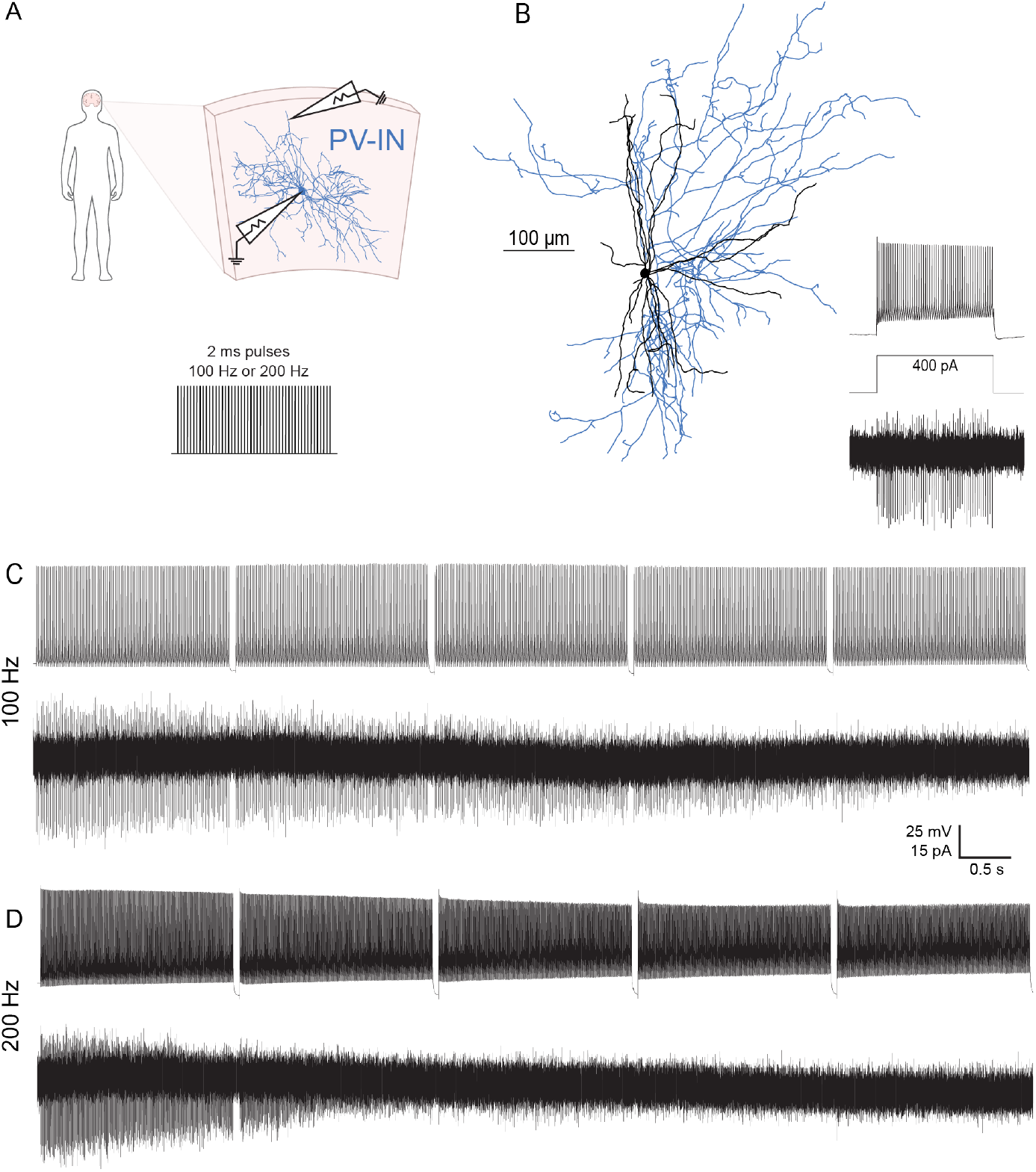
Human fast spiking INs display use-dependent changes in the axonal AP waveform during brief depolarizing pulses. (A) Experimental schematic for paired whole-cell somatic and axon-attached recordings in human neocortical fast-spiking INs during brief depolarizing pulses. (B) Morphologic reconstruction and example voltage trace in response to a square wave depolarization from a human fast-spiking aspiny presumptive basket cell. (C-D) Examples of simultaneous whole-cell somatic voltage (top) and axon-attached current (bottom) traces during brief depolarizing pulses delivered at the soma pipette at (C) 50 Hz and (D) 100 Hz.

### Axonal AP propagation is largely unchanged in neocortical PV-INs during physiologic AP trains

Because the response to long trains of repetitive brief depolarizing pulses and square wave current injections (Figs. S3, S2, 3) do not reflect PV-IN firing patterns *in vivo*, we sought to assess axonal AP propagation during more physiologic PVIN firing patterns. We performed simultaneous whole-cell somatic and distal axon-attached recordings during a 5-min pulse train simulating the in vivo firing pattern of an optotagged PV-IN from the publicly available Allen Institute Behavior Neuropixels dataset (Fig. 4). Throughout the selected 5-minute window, the cell exhibited bursts of firing that averaged over 50 Hz for several seconds, achieving instantaneous firing frequencies of up to 200 Hz for brief periods. During this simulated physiologic activity regime, there was not significant modulation of the axonal AP waveform in PV-INs, and thus conduction remained robust (Fig. 4B-D).

**Fig. 4.**
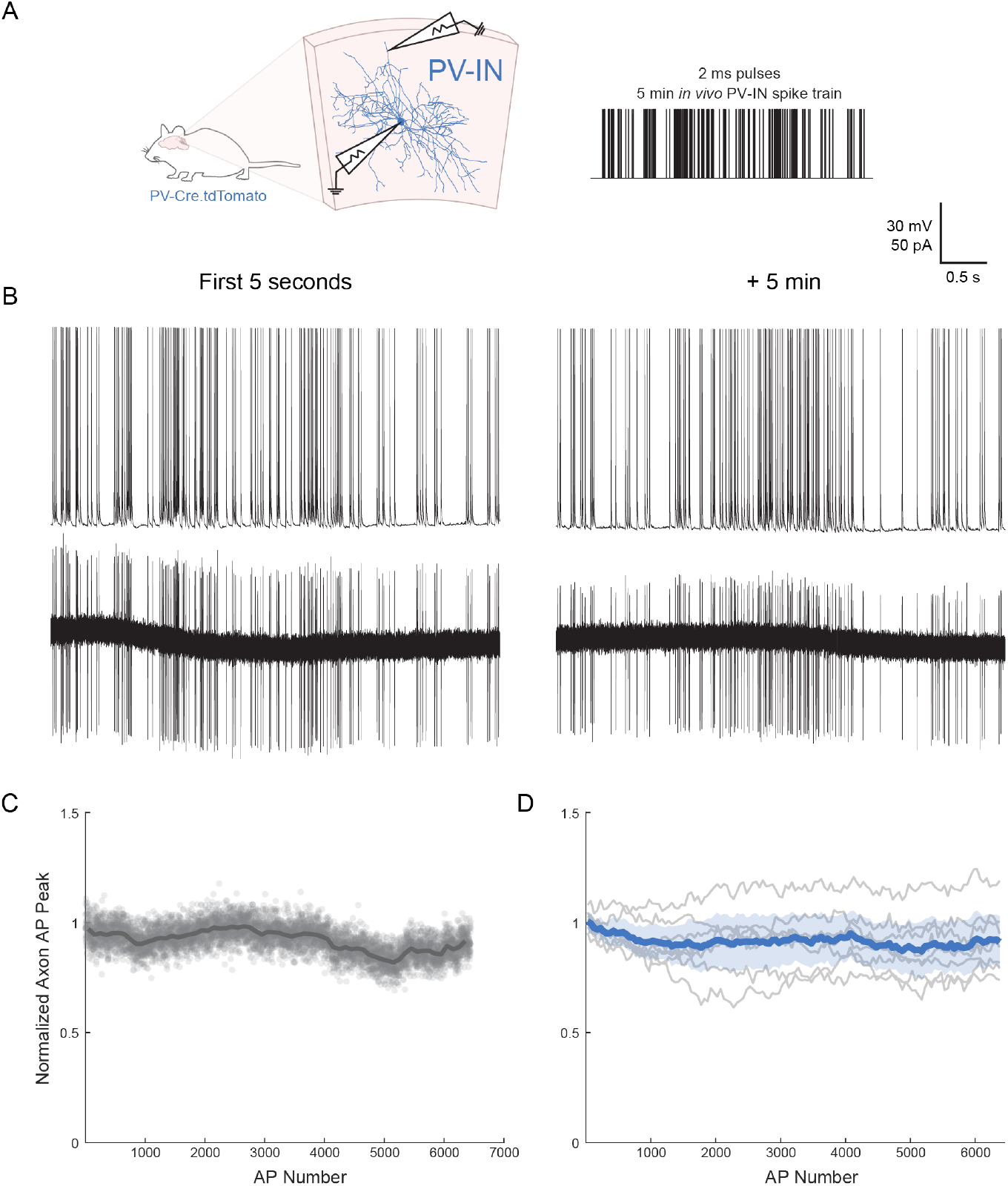
Neocortical PV-IN axonal AP propagation fidelity remains high during physiologic AP trains. (A) Experimental schematic of paired whole-cell somatic and axon-attached recordings in PV-INs (left) during delivery of a 5 min pulse train simulating *in vivo* PV-IN firing at the soma. (B) Example simultaneous whole-cell somatic voltage (top) and axon-attached current (bottom) traces during a physiologic AP train (right). (C-D) Axon AP peak (normalized to the mean of the first 5 AP peaks) vs. AP number in (C) an example cell and (D) as a summary across cells (dark blue indicates the mean, shaded region indicates the 95% confidence intervals; *n* = 8 cells from *N* = 4 mice).

### PV-INs display decreases in axonal AP propagation fidelity during seizure-like events

In contrast to physiologic activity in which neocortical PV-INs exhibit high frequency firing (>100 Hz) for only short bursts, PV-INs can fire at high frequency for several seconds preceding and during the onset of focal seizures (21). These long epochs of high-frequency firing occur in the setting of synchronous activity across nearby neurons, leading to metabolic and ion concentration derangements that may impact PV-IN axonal AP propagation (37–42). In an attempt to assess PV-IN axonal AP propagation during pathologic activity regimes at the onset of focal seizures, we performed whole-cell somatic and axon-attached recordings of neocortical PV-INs in the well-established bicuculline-induced model of seizure-like events in the acute brain slice (Fig. 5) (43, 44).

**Fig. 5.**
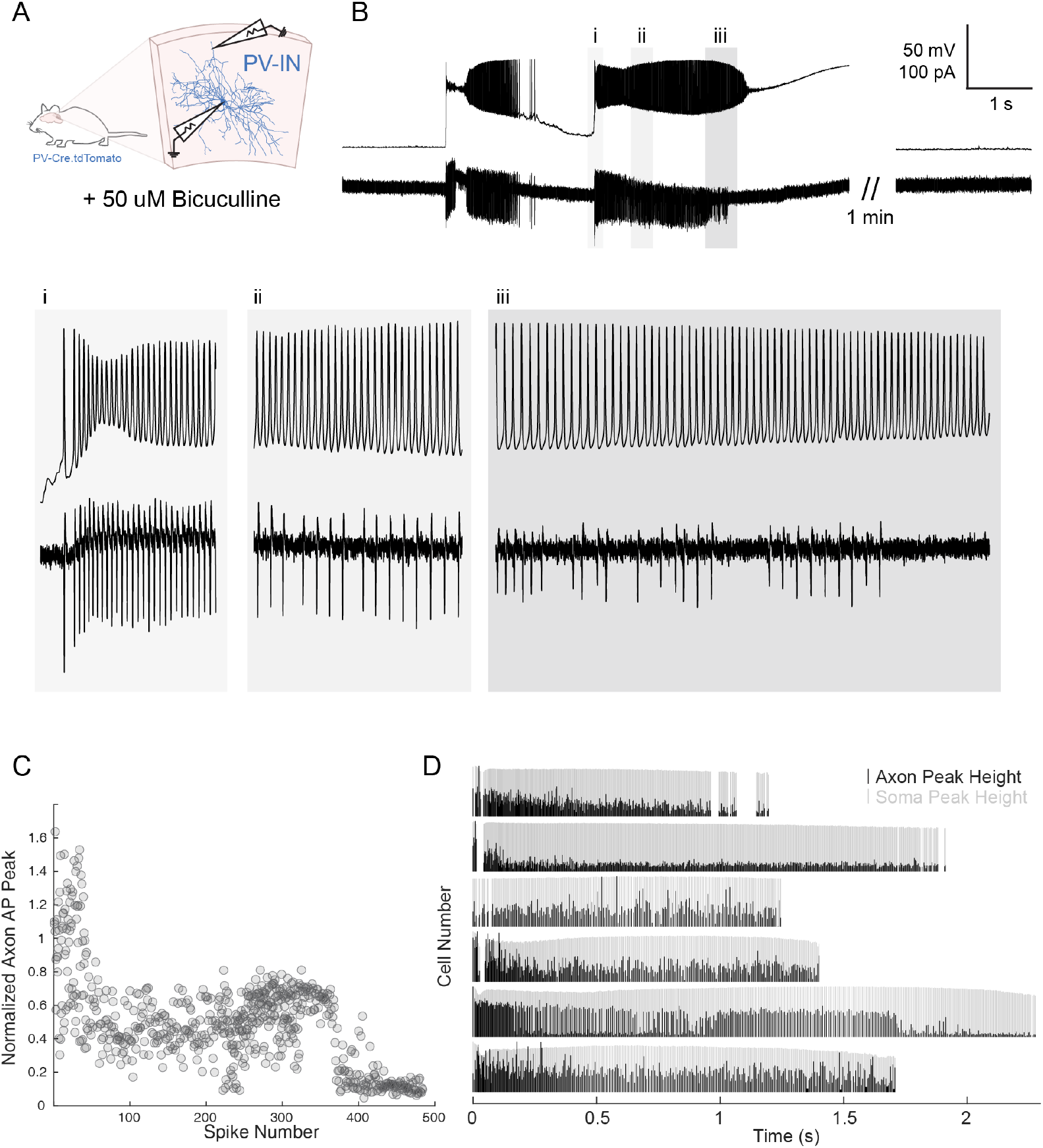
PV-INs display progressive decreases in axonal AP propagation fidelity during seizure-like events induced by 50 µM bicuculline. (A) Experimental schematic for paired whole-cell somatic and axon-attached recordings in neocortical PV-INs during seizure-like events induced by 50 µM bicuculline. (B) Example of a simultaneous whole-cell somatic voltage (top) and axon-attached current (bottom) trace from a PV-IN during spontaneous discharges within a seizure-like event. (C) Plot of axon AP peak (normalized to the mean of the first 5 AP peaks) vs. AP number during the final spontaneous discharge during the seizure-like event. (D) Raster plot of APs over time during the final spontaneous discharge of the seizure-like event. Black lines indicate the height of axon AP peaks (normalized to the mean of the first 5 axon AP peaks). Gray lines indicate the height of soma AP peaks (normalized to the height of the max soma AP peak) (*n* = 6 cells from *N* = 4 mice).

Following bath application of 50 µM bicuculline, recorded cells exhibited spontaneous repetitive (2-6) epileptiform-like discharges during which cells fired hundreds of spikes at frequencies of up to 400 Hz (Fig. 5B-D). We observed a striking dissociation between the AP recorded at the soma and the AP recorded at the distal axon during these events. Early in the discharge, some soma/AIS-generated APs only produced spikelets or low-amplitude APs at the soma but were recorded as currents reflective of full spike height at the more distal axon recording site (a phenomenon which has been reported previously in neocortical pyramidal cells during picrotoxin-induced seizure-like events) (45). In contrast, APs occurring later in the epileptiform-like discharge led to progressively smaller or undetectable currents in the distal axon (Fig. 5B-D). These results suggest that, during seizure-like events, there is a decrease in AP propagation fidelity in neocortical PV-INs. This phenomenon may be relevant to the pathophysiology of focal-onset seizures and other states associated with high PV-IN firing frequency.

### PV-IN synaptic boutons show use-dependent decreases in Ca^2+^ concentration and synaptic transmission

Next, we sought to understand how use-dependent decreases in axonal AP propagation in PV-INs affect spike-evoked bouton Ca^2+^ influx and associated neurotransmitter release (46). To assess Ca^2+^ dynamics at neocortical PVIN axonal boutons, we patched and filled neocortical PV-INs with the synthetic Ca^2+^ indicator Fluo-5F and imaged *en passant* presynaptic boutons during trains of APs at 100 Hz (Fig. 6). Consistent with previous recordings of presynaptic bouton Ca^2+^ dynamics during short trains of APs, we observed summation of the Fluo-5F fluorescence signal early in the AP train (Fig. 6B-C) (8, 47–49). However, after the cell had generated ∼600 APs, we observed a steady decrease in the Fluo-5F signal. This finding is consistent with a progressive decrease in the amplitude and/or success rate of APs invading the bouton during prolonged, high-frequency activity. In line with the progressive decrease of the Fluo-5F Ca^2+^ signal, we observed decreases in the amplitude and success rate of PVIN-mediated synaptic transmission onto layer 2-3 pyramidal cells during prolonged trains of APs (Fig. S5).

**Fig. 6.**
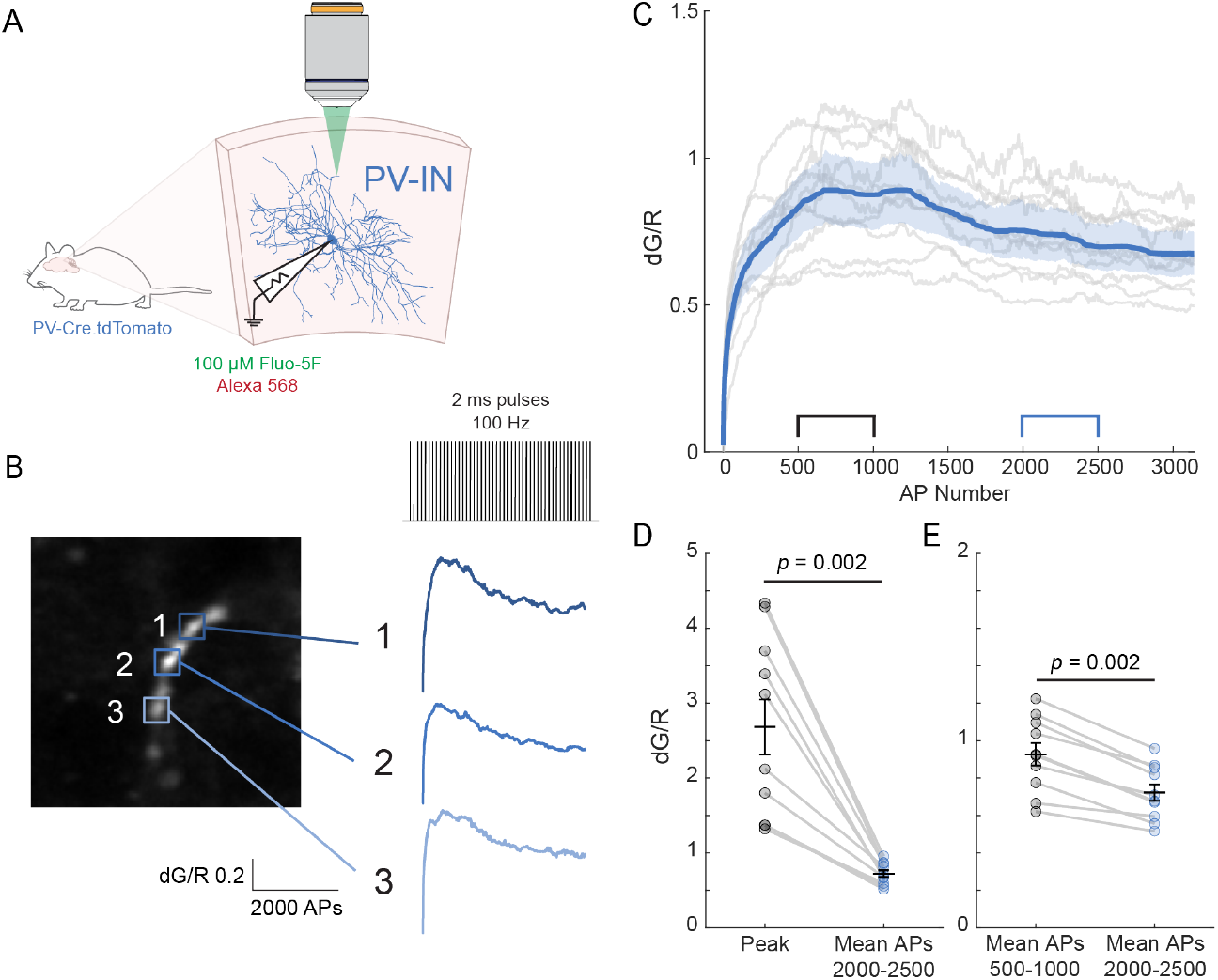
Use-dependent decreases in Ca2+ signal in PV-IN boutons. (A) Experimental schematic for recordings of Fluo-5F imaging in neocortical PV-IN boutons during brief depolarizing pulses delivered at the soma. (B) Example image and Ca^2+^ traces from *en passant* boutons in the distal axon of a PV-IN. (C) Summary data of dG/R (change in green Fluo-5F fluorescence normalized to change in red fluorescence from non-functional fluorophore) vs. AP number during brief depolarizing pulses at 100 Hz (10 boutons from *n* = 5 cells from *N* = 3 mice). Light gray lines indicate traces from individual cells (with median filter smoothing), dark blue line indicates the mean across cells, light blue shaded region indicates 95% confidence intervals. (D) Quantification of dG/R values from (C) showing (D) peak dG/R values vs. mean across APs 2000-2500 and (E) mean across APs 500-1000 vs. mean across APs 2000-2500. Bars represent mean and standard error of the mean. *P*-values calculated using the Wilcoxon signed-rank test.

### A model based on a sum of exponential decays describes use-dependent regulation of axonal AP propagation fidelity

To generate hypotheses as to which biological processes might underlie use-dependent changes in the axonal AP waveform in PV-INs, we developed a mathematical model of the data. Because we observed a progressive decrease in the axon AP peak with each action potential, accompanied by a time-dependent “recovery” in the axonal AP peak between APs, we fit our data to a summation of exponential decays model (Fig. 7, see Materials and Methods). To ensure applicability across multiple firing regimes, we fit data from 3 separate protocols: regular brief depolarizing pulses (100 and 200 Hz), brief depolarizing pulses at varying frequencies (200, 100, and 50 Hz), and square wave depolarizations (600 ms current step every 3 seconds, starting at −100 pA and increasing by 25 pA steps) (Fig. 7A-D). We observed a robust fit to this model across cells and protocols, with a median R^2^ value of 0.76 (Fig. 7E). The median value for the decay fitting parameter (*τ*) was 11.23 sec, which suggests that previous APs impact subsequent axonal AP propagation on a timescale on the order of seconds. Thus, significant decrements in axonal APs are observed within a window following prolonged trains of APs sufficient to engage this underlying use-dependent mechanism yet before appearance of a timedependent recovery process.

**Fig. 7.**
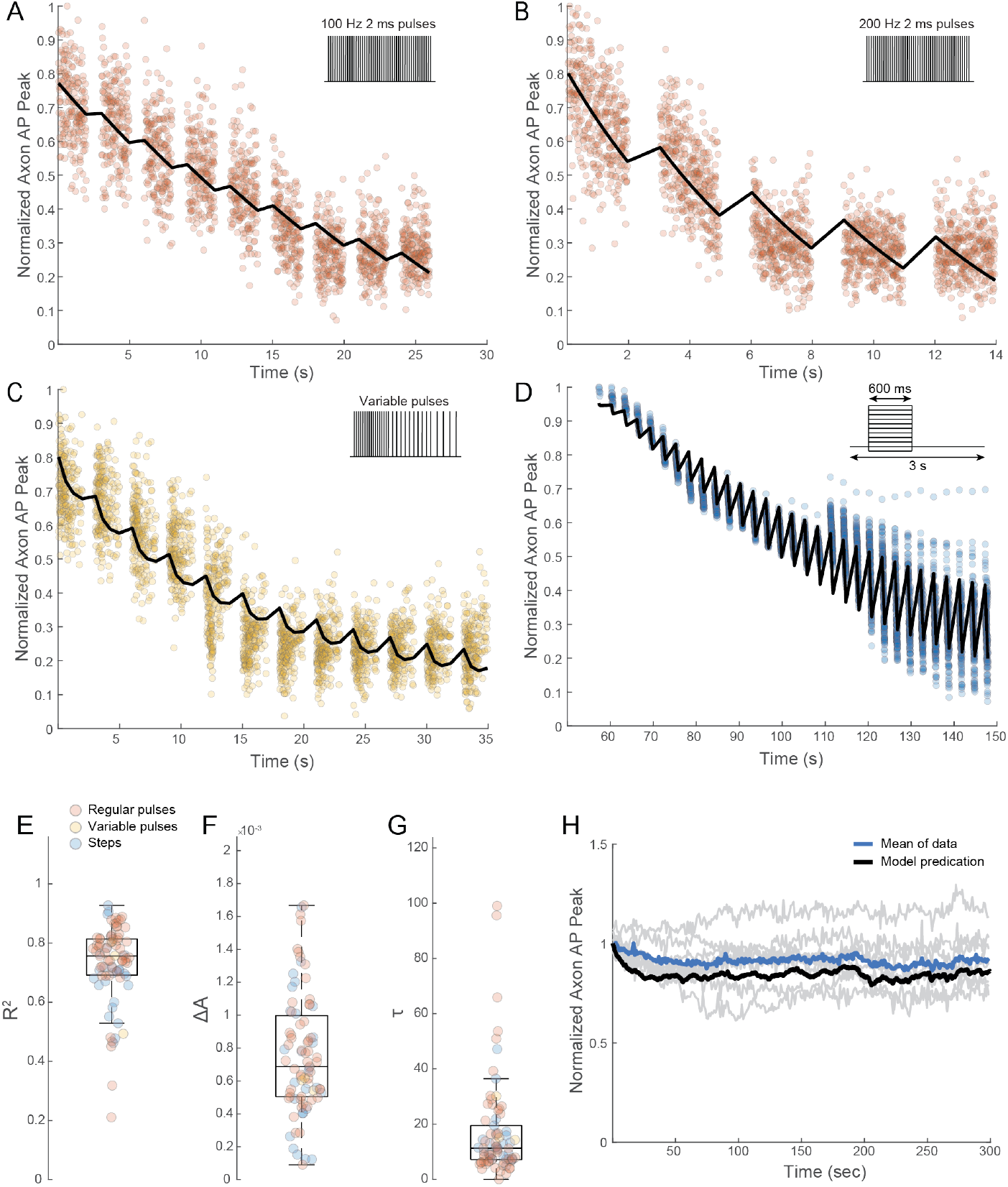
Use-dependent regulation of the axonal AP waveform can be modeled as a sum of exponential decays. (A-D) Example plots of raw normalized axon AP peaks vs. time data for cells stimulated with (A) brief pulses at 100 Hz, (B) brief pulses at 200 Hz, (C) pulses at variable frequencies, and (D) square wave depolarizations. (E-G) Box and whisker plots of the (E) goodness of fit R^2^ (median = 0.76), (F) ∆A fitting parameter (median = 0.69e10^-3^), and (G) decay (*τ*) fitting parameter (median = 11.23 s) across cells. Data points colored by stimulation protocol (83 recordings from *n* = 52 cells from *N* = 35 mice). (H) Plot of axon AP peaks (normalized to the mean of the first 5 APs) vs. time for the simulated *in vivo* spike train data in **Figure 4** (gray lines indicate individual cells, blue line indicates mean across cells) overlaid with axonal AP peaks predicted by the sum of exponential decays model parameterized with the median values from (F) and (G) (with outliers excluded).

### High extracellular K^+^ can mimic use-dependent regulation of axonal fidelity

The values for the decay (*τ*) fitting parameter from our sum of exponential decays model (Fig. 7) suggested that the underlying mechanism of use-dependent decrements in the axonal AP is a biological process (or multiple biological processes) operating on the order of seconds. One candidate mechanism is decreased excitability of the axon due to use-dependent accumulation of periaxonal K^+^, leading to depolarization of the axonal membrane potential and thus decreased Na^+^ channel availability (due to Na^+^ channel inactivation) (50–53), which would explain the observed use-dependent decrease in axonal AP height. To determine if modifying extracellular K^+^ concentration affects PV-IN axonal AP propagation, we performed paired wholecell somatic and axon-attached recordings in neocortical PVINs in standard ACSF containing 2.5 mM K^+^, then washed on ACSF containing 10 mM K^+^ (Fig. 8). As expected, 10 mM K^+^ ACSF led to a depolarization of the somatic membrane potential, which we offset with a DC current injection to hold the soma at −70 mV to control for the effect of resting membrane potential on the somatic AP waveform. Wash-on of ACSF with 10 mM K^+^ resulted in a marked decrease in the axonal AP signal, without a change in the peak of the somatic AP (Fig. 8B-E). This effect was partially abrogated by re-wash-on of ACSF with 2.5 mM K^+^ in a subset of cells (Fig. 8B,E). These findings suggest that extracellular K^+^ concentration can serve as a modulating factor of AP propagation in PV-IN axons.

**Fig. 8.**
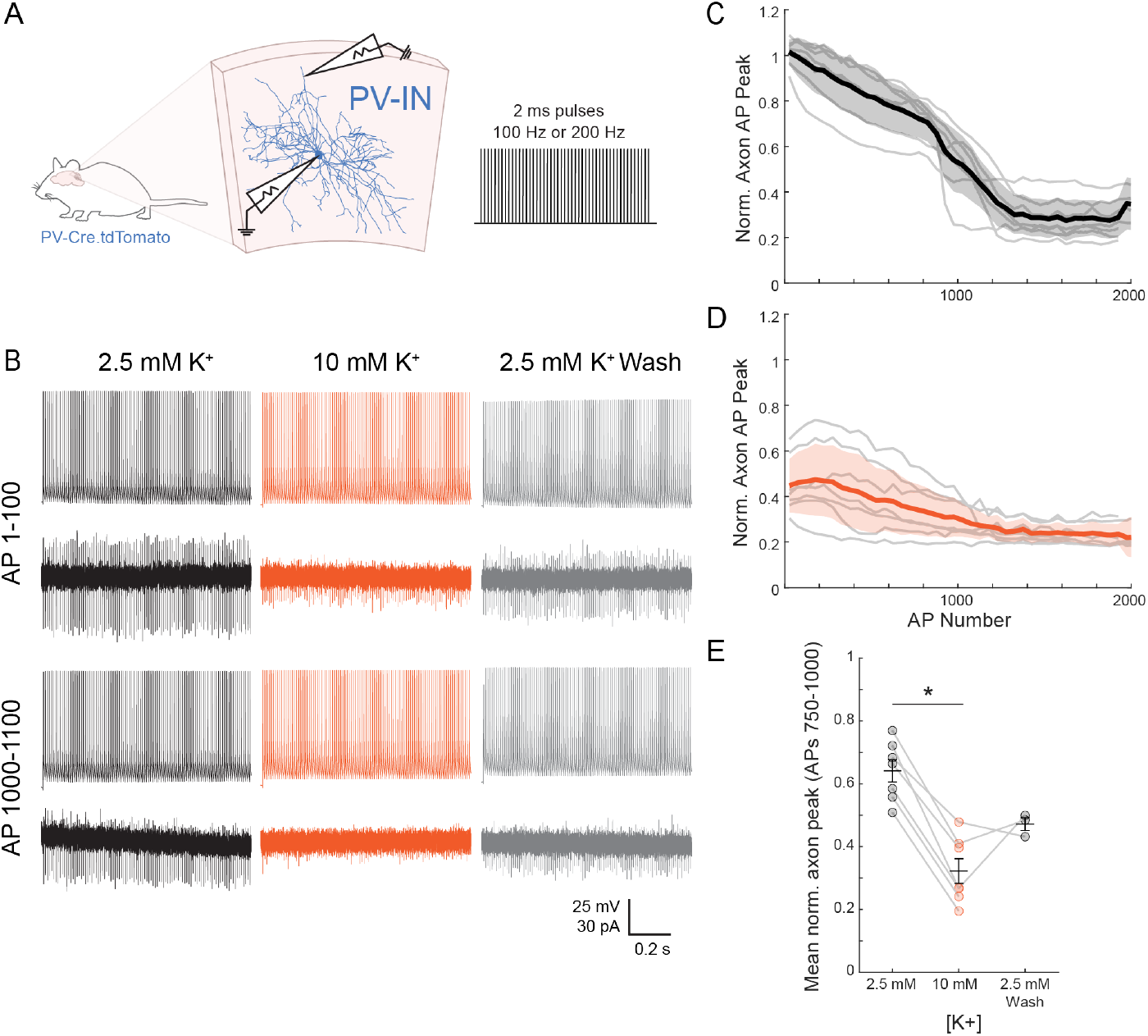
High extracellular K+ can mimic use-dependent modulation of the axonal AP. (A) Experimental schematic for paired whole-cell somatic and axon-attached recordings in neocortical PV-INs during brief depolarizing pulses while changing the concentration of extracellular K^+^. (B) Examples of simultaneous whole-cell somatic voltage (top) and axon-attached current (bottom) traces during brief depolarizing pulses delivered at the soma pipette at 100 Hz in ACSF with 2.5 mM K^+^, after wash-on of ACSF with 10 mM K^+^, and again after wash-on of 2.5 mM K^+^. (C-D) Summary data of axonal AP peak (normalized to the mean of the first 5 APs) vs. AP number during 100 Hz brief depolarizing pulses (C) at baseline in ACSF with 2.5 mM K^+^ and (D) after wash-on of ACSF with 10 mM K^+^. Gray lines indicate individual cells, while the dark black/orange line indicates the mean across cells with the shaded blue region indicating standard error of the mean (*n* = 7 cells from *N* = 4 mice). (E) Summary data of mean normalized axon AP peak for APs 750-1000 in 100 Hz pulse train in ACSF with 2.5 mM K^+^, after wash-on of ACSF with 10 mM K^+^ (*p* = 0.0156, Wilcoxon signed-rank test), and again after wash-on of 2.5 mM K^+^.

## Discussion

Here we show that, in line with previous studies, PV-IN axonal spike propagation is robust during brief periods of moderate firing frequency (Fig. 4). However, in PV-INs from both mouse and human, there are significant decreases in the axonal AP waveform during prolonged, high-frequency firing of various forms, even while somatic AP generation remains stable (Figs. 2, 3, S2, 5), demonstrating soma:axon dissociation in the effect of repetitive firing on AP waveform. Our data further suggests that, at least via the methods employed here, the status of axonal AP propagation cannot be clearly assigned a binary “success” or “failure.” Additionally, usedependent changes in the axonal AP waveform can be modeled as a sum of exponential decays (Fig. 7) with a decay time constant on the order of seconds, suggesting a relatively slow biological process(es), such as derangements of ion concentration gradients during prolonged, high-frequency activity (Fig. 8).

### Mechanisms of use-dependent decreases in axonal AP fidelity

We chose to conduct our axon-attached recordings in voltage-clamp mode, given that the patch (which can be modeled as an RC circuit) acts as a low-pass filter, thus distorting the amplitude and kinetics of a fast event like an AP (27).The AP-associated current measured in axonattached voltage clamp is largely capacitive because the impedance across the capacitance of the patch is lower than the impedance across the resistance of the patch during fast events (27, 29). There are several factors that affect the amplitude of the currents isolated during axon-attached recordings in voltage clamp: (1) the magnitude of the change in axonal resting membrane potential (.6.V_m_) during the AP (with a larger .6.V_m_ resulting in larger currents), (2) the rise time of the axonal AP (with a faster rise time leading to a larger current), (3) the resistance of the seal (R_seal_) relative to the electrode (R_elec_) (with a larger R_seal_/R_elec_ leading to a larger current), and (4) the capacitance of the patch (C_patch_) (with a larger C_patch_ leading to a larger current). Assuming no change in R_seal_, R_elec_, or in C_patch_ over the course of the recording, use-dependent decreases in the amplitude of the axonal AP-driven currents must be due to (1) decreases in .6.V_m_ during the AP and/or (2) increases in the rise time of the AP (i.e., a shorter and/or slower AP). A decrease in .6.V_m_ could be due to either a more depolarized resting V_m_ from which the AP departs and/or a decrease in the peak voltage of the AP. Hence, there are three main changes in the axonal AP voltage waveform that could underlie the use-dependent decreases in the amplitude of the axonal AP currents observed here: progressive depolarization of the V_m_ of the distal axon, progressive decreases in the peak voltage of the axonal AP, or a progressive increase in the rise time of the axonal AP.

There are multiple mechanisms that could result in these changes in the axonal AP voltage waveform. Progressive depolarization of axon V_m_ could be due to changes in the ion concentration gradients across the axonal membrane. In small structures such as neocortical PV-INs axons (with an average diameter ∼0.3 µm), the classic cable theory assumption that bulk ion concentrations remain constant during activity may not hold (54, 55). Prolonged high-frequency activity will lead to extracellular accumulation of K^+^ ions and/or intracellular accumulation of Na^+^ ions if activity outpaces the restoration of ion concentration gradients by the axonal Na^+^-K^+^ ATPase and/or astrocytic clearance mechanisms (55–57). In support of this possible mechanism, there is extensive evidence of increases in extracellular K^+^ during high-frequency activity in the mammalian cerebral cortex (40, 41, 58, 59). It is also possible that increases in axonal V_m_ could be caused by slow activation of a depolarizing conductance. For example, recent work has demonstrated expression of hyperpolarization-activated cyclic nucleotide-gated channels in PV-IN axons, which opposes hyperpolarization by the axonal Na^+^-K^+^ ATPase (11).

The other two possible changes in the axonal AP voltage waveform – namely, progressive decreases in the voltage peak or increases in the rise time of the axonal AP – likely result from decreases in the availability of axonal voltagegated Na^+^ channels (60). Decreases in Na^+^ channel availability could occur as a secondary consequence of a progressive depolarization of the axonal V_m_ and/or slow recovery from inactivation of distal axonal Na^+^ channels. Studies across species and cell types have demonstrated that the kinetics of Na^+^ channel recovery from inactivation have a non-linear dependence on previous activity, with time constants reaching several seconds after prolonged, high-frequency firing (61– 64). Progressive decreases in Na^+^ channel availability could also be due to progressive activation or slow recovery from inactivation of distal axonal K^+^ channels (either hyperpolarizing the axon or preventing recovery of Na^+^ channels from inactivation). Despite the fact that Kv3.1/Kv3.2 channels (the dominant K^+^ channels in PV-IN axons) are non-inactivating on short time scales, Kv3.1 channels exhibit slow accumulation of inactivation on the order of seconds, particularly during brief repetitive depolarizations (65–69). Interestingly, this slow “U-type” inactivation is exacerbated under conditions of increased extracellular K^+^, which may be relevant during periods of high cell-intrinsic and/or network activity leading to extracellular K^+^ accumulation (68).

### Consequences of use-dependent regulation of axonal AP propagation

How does use-dependent modulation of the axonal AP waveform interact with the kinetics of synaptic depression in PV-INs? One possibility is that this phenomenon underlies, at least in part, the slow synaptic depression at PV-IN inhibitory synapses during prolonged activity. Previous work in fact suggests that a mechanism of synaptic depression in basket cells in visual cortex and dentate gyrus lies upstream of the exocytic step (70, 71). Progressive reductions in the amplitude of the axonal AP could lead to shorter duration APs and thus reductions in spike-evoked bouton Ca^2+^ influx and result in an apparent enhancement of synaptic depression (8, 72, 73). Another possibility is that use-dependent decreases in the axonal AP waveform could actually be serving to resist synaptic depression by preventing inactivation of bouton Ca^2+^ channels or depletion of the readily-releasable vesicle pool (74). This could ultimately serve to control synaptic depression and maintain some synaptic efficacy during prolonged high-frequency activity. These mechanisms could also be acting across multiple timescales. For example, use-dependent decrements in the axonal AP waveform could serve to resist synaptic depression during brief periods of activity, but ultimately exacerbate synaptic depression during periods of prolonged and/or high-frequency activity.

What are the consequences of use-dependent decreases in PV-IN AP propagation on circuit function? Prolonged, highfrequency PV-IN activity concordant with the stimulation paradigms employed in this study has been reported leading up to and at the onset of seizures in recordings from both animal models of epilepsy and human patients with focal-onset seizures (21, 24, 75). Similarly, in the bicuculline slice model of seizure-like events, we directly observed profound decreases in axonal AP propagation and clear failures during epileptiform-like discharges (Fig. 5). As such, it is possible that increased activity of PV-INs at the onset of focal seizures or at the edge of a propagating seizure wave could serve to paradoxically worsen inhibition if the activity is high enough to drive use-dependent decreases in axonal AP propagation fidelity (76). In other words, “increased” recruitment of inhibition by PV-INs during focal seizure onset and/or propagation may actually serve to decrease inhibitory output onto excitatory cells.

### Limitations of this study

The small caliber of neocortical IN axons renders whole-cell patch clamp recordings from single intact, uncut axons technically challenging or impossible (vs. “blebs,” in which the underlying actin cytoskeleton of the axon is disrupted, which may influence ion channel clustering) (77). As such, this study was limited to axon-attached recordings in the acute brain slice model. The acute brain slice cannot completely replicate normal glial function, depolarized membrane potential, presence of neuromodulators, and/or ongoing synaptic activity present in intact circuits *in vivo* (78, 79). Technical advances may facilitate whole-cell recordings or voltage imaging of axons *in vitro* and *in vivo*, which would allow for assessment of axonal AP propagation across behavioral and disease states. New tools for axon-specific manipulations (such as opsins with highly-specific axon targeting and/or increased spatial precision of photostimulation) could facilitate causal interrogation of the effects of decreases in axonal AP propagation (80). For example, the above hypothesis that inhibition of axonal AP propagation fidelity in nearby PV-INs exacerbates inhibitory restraint during focal seizures *in vivo* could be tested directly. In summary, this study demonstrates a dissociation between somatic AP generation and the axonal AP in PV-INs, with relevance for normal physiological function as well as neurological disorders such as epilepsy. As such, the assumption that all APs recorded at the soma, such as those recorded *in vivo* using extracellular electrophysiology or Ca^2+^ imaging, successfully propagate down the axon in a binary manner to reliably invade pre-synaptic terminals and trigger neurotransmitter release, may not hold under all conditions.

## ACKNOWLEDGEMENTS

We thank Emily Hoddeson and Xiaohong Zhang for technical assistance with mouse colony management and Benjamin C. Kennedy for access to surgical specimens.

## Funding

National Institutes of Health Grant F31NS132519 (SRL) National Institutes of Health Grant R01NS119977 (EMG)

## Author contributions

Conceptualization: SRL, EMG; Methodology: SRL, EMG; Investigation: SRL; Visualization: SRL; Supervision: EMG; Writing—original draft: SRL; Writing—review & editing: SRL, EMG

## Competing interests

Authors declare that they have no competing interests.

## Data and materials availability

All data are available in the main text or the supplementary materials. All original code has been deposited at https://github.com/GoldbergNeuroLab/LiebergallGoldberg-2025. Any additional information required to reanalyze the data reported in this paper is available from the lead contact upon request. This study did not generate new unique materials.

## Supplementary Figures

**Fig. S1.**
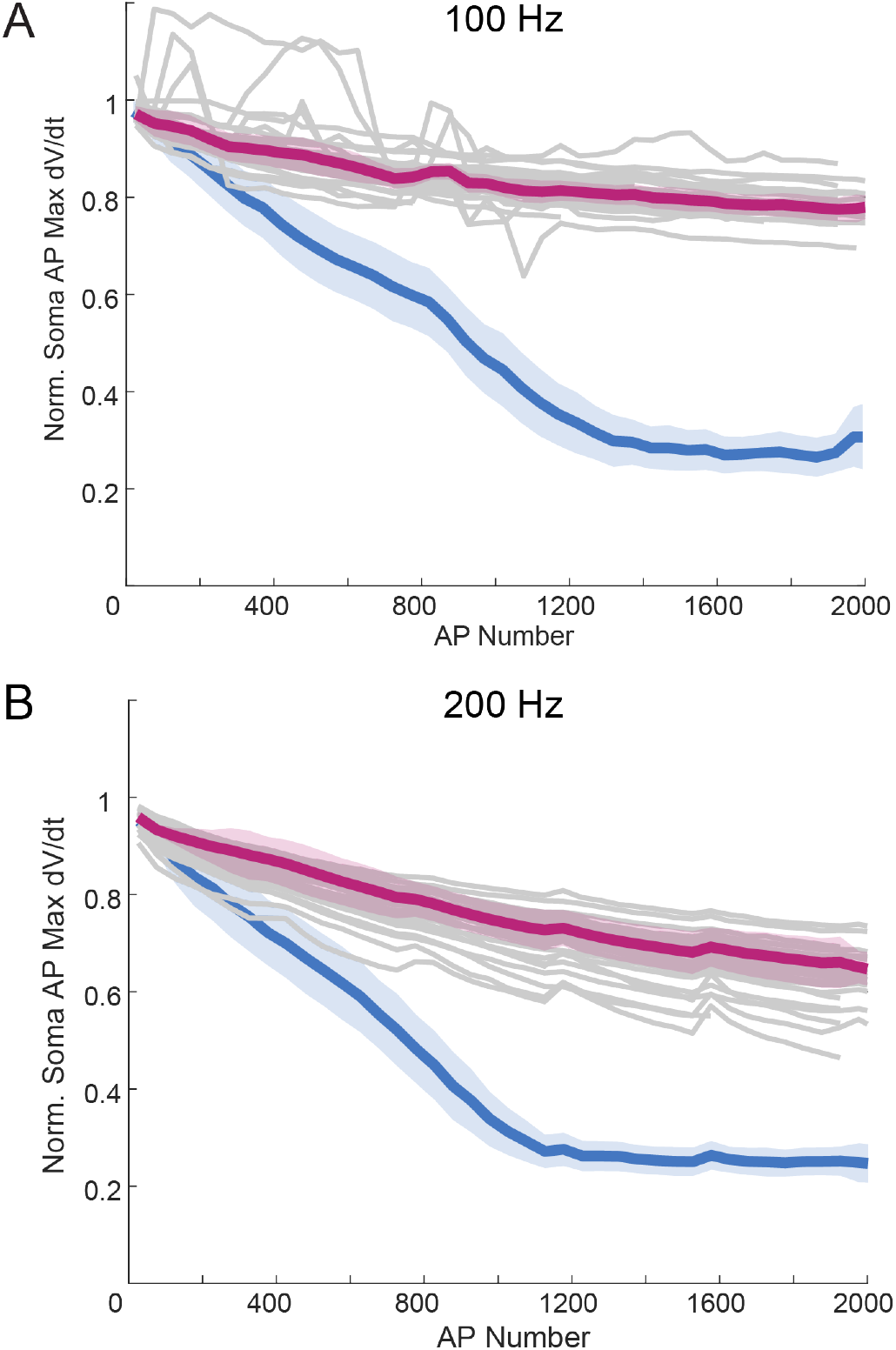
Use-dependent changes in PV-IN axonal AP waveforms outpace use-dependent changes in the somatic AP max dV/dt. (A-B) Summary data of somatic AP max dV/dt (normalized to the mean of the first 5 APs) vs. AP number at (A) 100 Hz and (B) 200 Hz. Gray lines indicate individual cells, the dark magenta line indicates the mean across cells, and the shaded magenta region indicates 95% confidence intervals (*n* = 24 cells from *N* = 14 mice). Dark blue lines and shaded regions are a replication of the data presented in Fig. 2F,H for visual comparison.

**Fig. S2.**
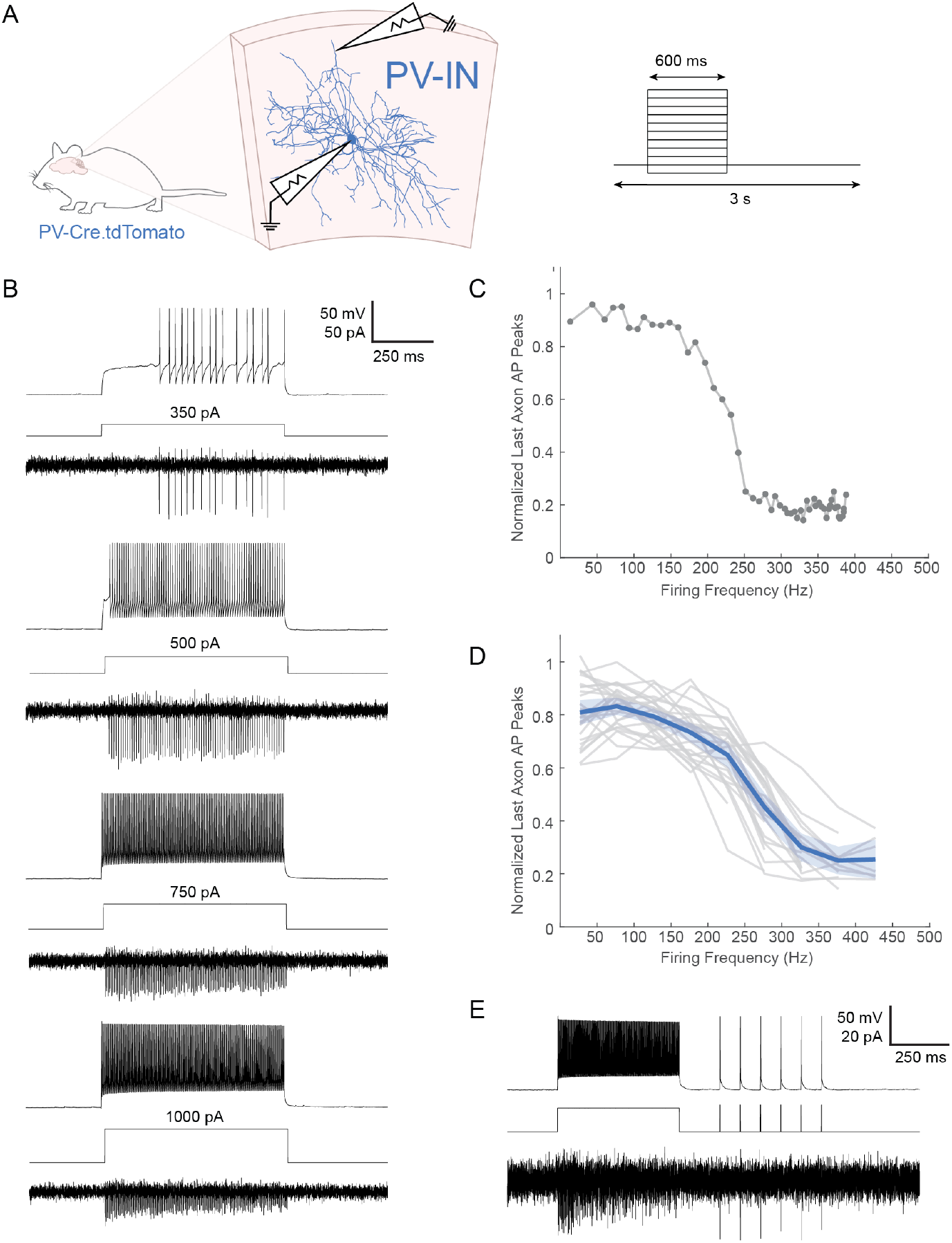
PV-INs display use-dependent changes in the axonal AP waveform fidelity during square wave depolarizations. (A) Experimental schematic for paired whole-cell somatic and axon-attached recordings in neocortical PV-INs during increasing square wave depolarizations. (B) Examples of simultaneous whole-cell somatic voltage (top) and axon-attached current (bottom) traces during square wave depolarizing pulses delivered at the soma pipette. (C-D) Plot of the mean of the last 5 axonal AP peaks in the current step (normalized to the mean of the first 5 APs in the recording) vs. firing frequency during the current step (C) for an example cell and (D) across multiple cells (gray lines indicate individual cells, dark blue line indicates mean across cells, shaded blue region indicates 95% confidence intervals; *n* = 33 cells from *N* = 23 mice). (E) Example trace of 20 Hz brief depolarizing pulses delivered after the current step, demonstrating that decreases in axonal AP peaks during current steps are not due to rundown in recording quality.

**Fig. S3.**
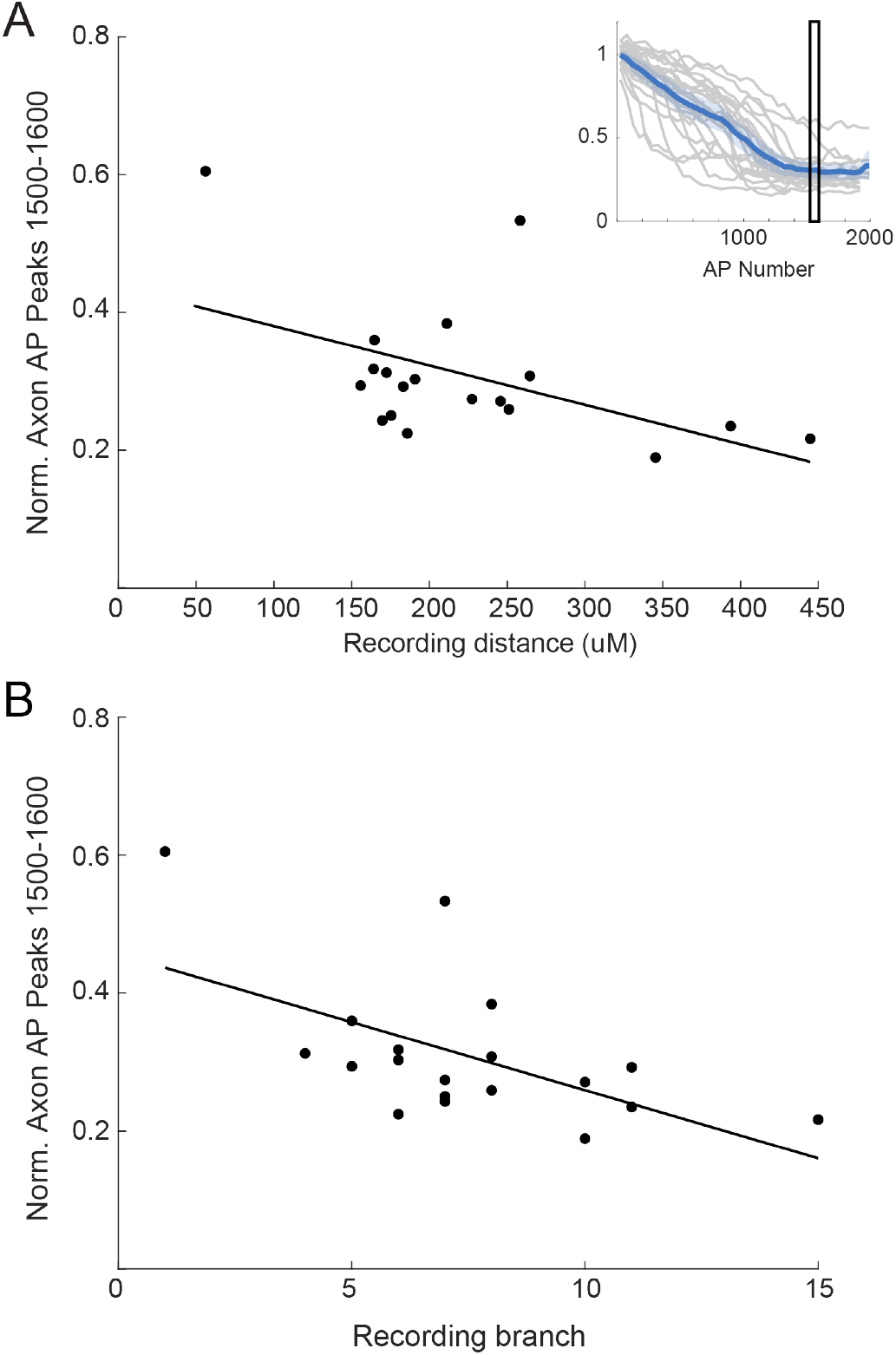
PV-IN axonal propagation depends on the distance and number of branch points traversed by the AP. (A) Plot of the mean of APs 1500-1600 (normalized to the mean of the first 5 APs in the recording) vs. the distance from the axon hillock to the recording site for the data in Figure 2F (*n* = 33 cells from *N* = 23 mice). Solid line indicates linear regression (R^2^ = 0.25, *p* = 0.03). (B) Plot of the mean of APs 1500-1600 (normalized to the mean of the first 5 APs in the recording) vs. branch order for the recording site. Solid line indicates linear regression (R^2^ = 0.34, *p* = 0.01).

**Fig. S4.**
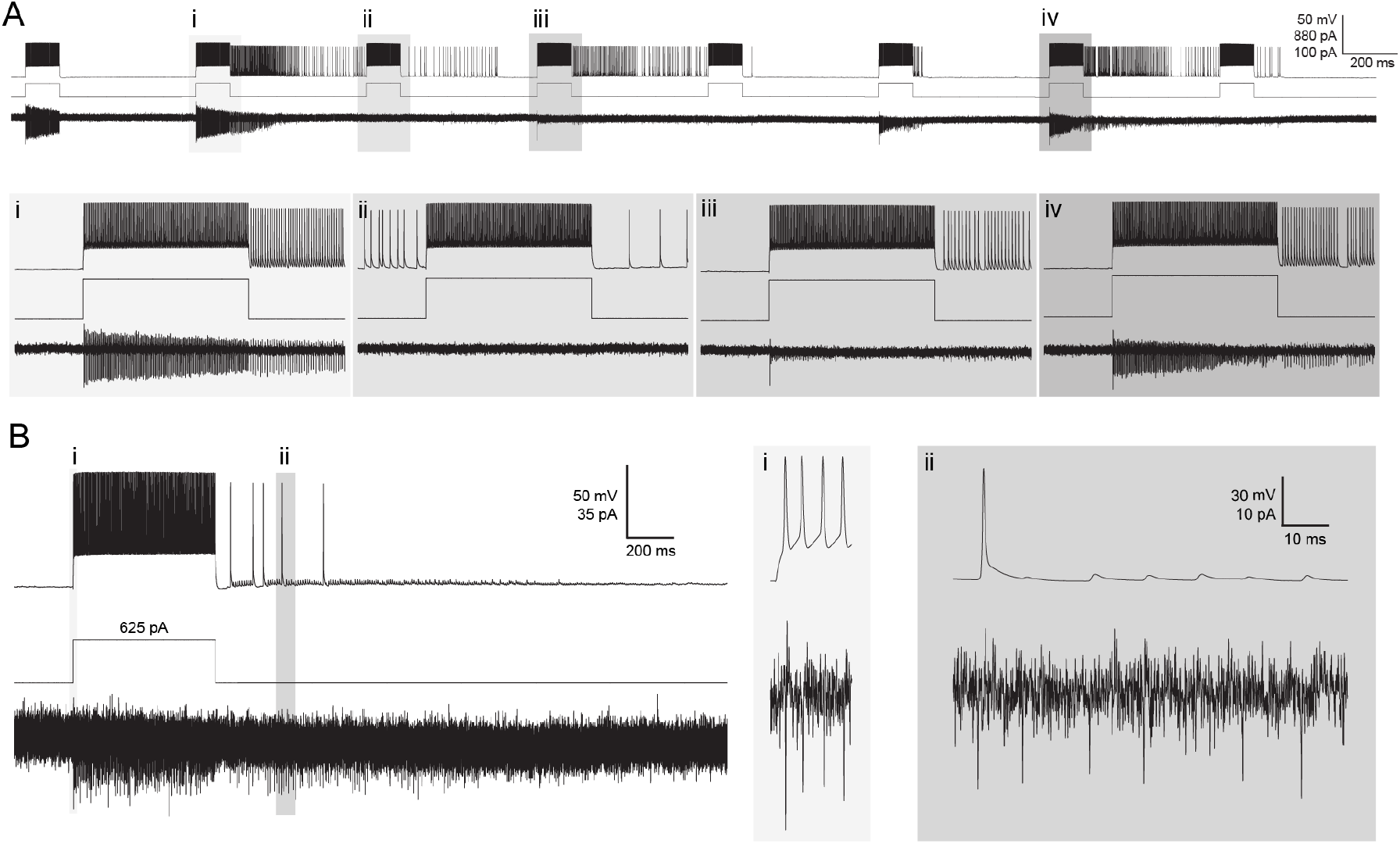
Barrages of ectopic APs are correlated with profound changes in the axonal AP waveform. (A) Example trace of a paired whole-cell somatic and axon-attached recording in a neocortical PV-IN in response to square wave depolarizations in which the cell displayed spontaneous persistent firing of APs generated in the distal axon. (B) Example trace in which axonal AP propagation fidelity of distal axon-generated APs is higher at the distal axon recording site relative to the somatic recording site.

**Fig. S5.**
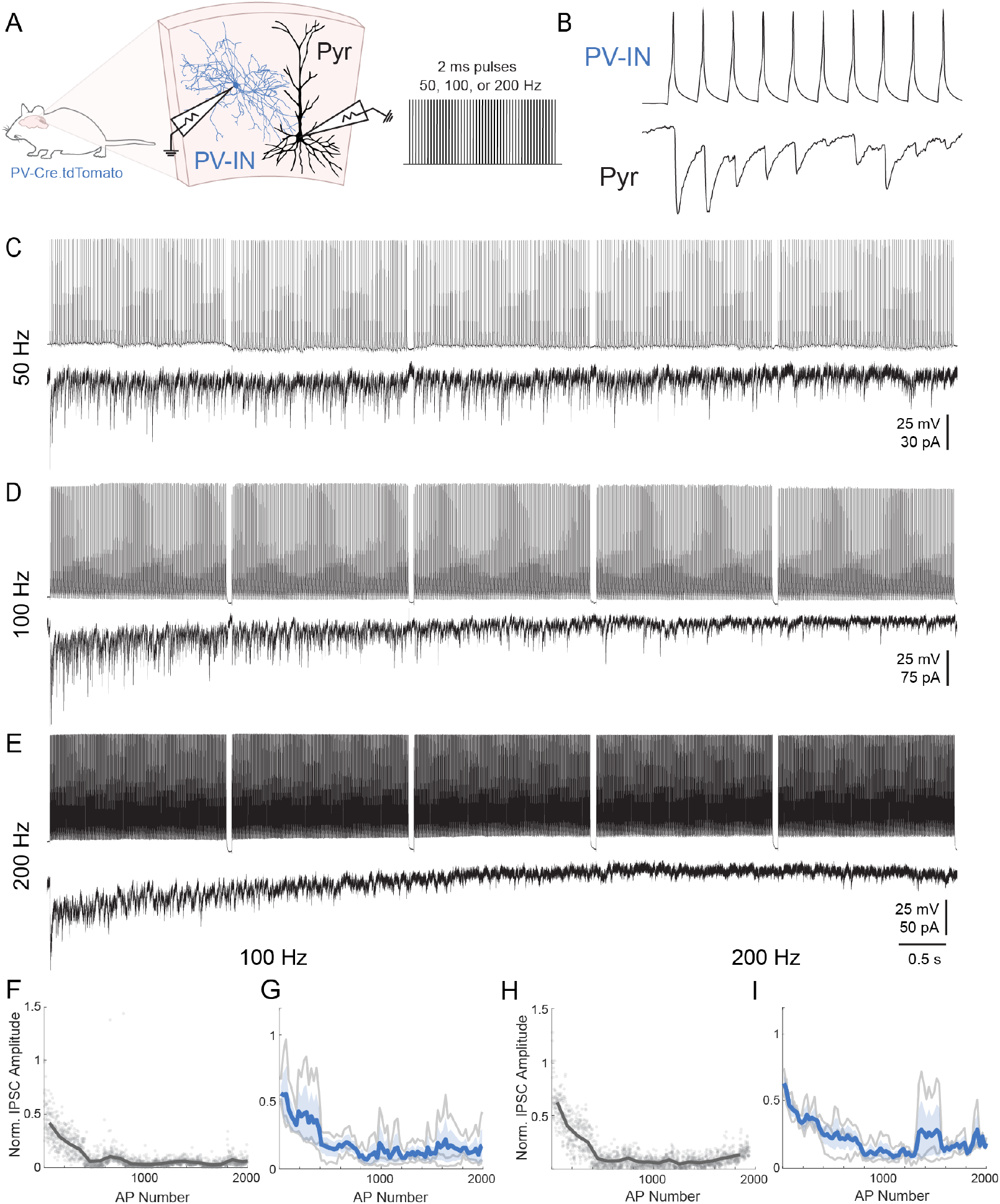
Use-dependent decreases in PV-IN-mediated synaptic transmission. (A) Experimental schematic for paired somatic recordings of synaptic transmission in a pre-synaptic PV-IN and a post-synaptic Layer 2/3 pyramidal neuron. Pre-synaptic PV-INs were stimulated with 2ms pulses at 50, 100, or 200 Hz. (B-E) Examples of simultaneous whole-cell somatic voltage traces from a presynaptic PV-IN (top) and current traces from a post-synaptic Layer 2/3 pyramidal cell (bottom) during brief depolarizing pulses delivered to the pre-synaptic cell at (B) 50 Hz, (C) 100 Hz, and (D) 200 Hz. (B) displays the first 10 APs from (D) at higher temporal resolution. (F,H) Plot of IPSC peak (normalized to the amplitude of the first evoked IPSC) vs. AP number for an example cell at (F) 100 Hz and (H) 200 Hz. Gray points indicate individual APs, while dark gray line indicates the mean across bins of 25 APs. (G,I) Summary data of IPSC peak (normalized to the amplitude of the first evoked IPSC) vs. AP number at (G) 100 Hz and (I) 200 Hz. Gray lines indicate individual cells, while the dark blue line indicates the mean across cells and the shaded blue region indicates standard error (*n* = 3 cells from *N* = 3 mice).

## Notes

### Competing Interest Statement

The authors have declared no competing interest.

